## Supplementary figures and images for "Omitting age-dependent mosquito mortality in malaria models underestimates the effectiveness of insecticide-treated nets"

### S1 Fig

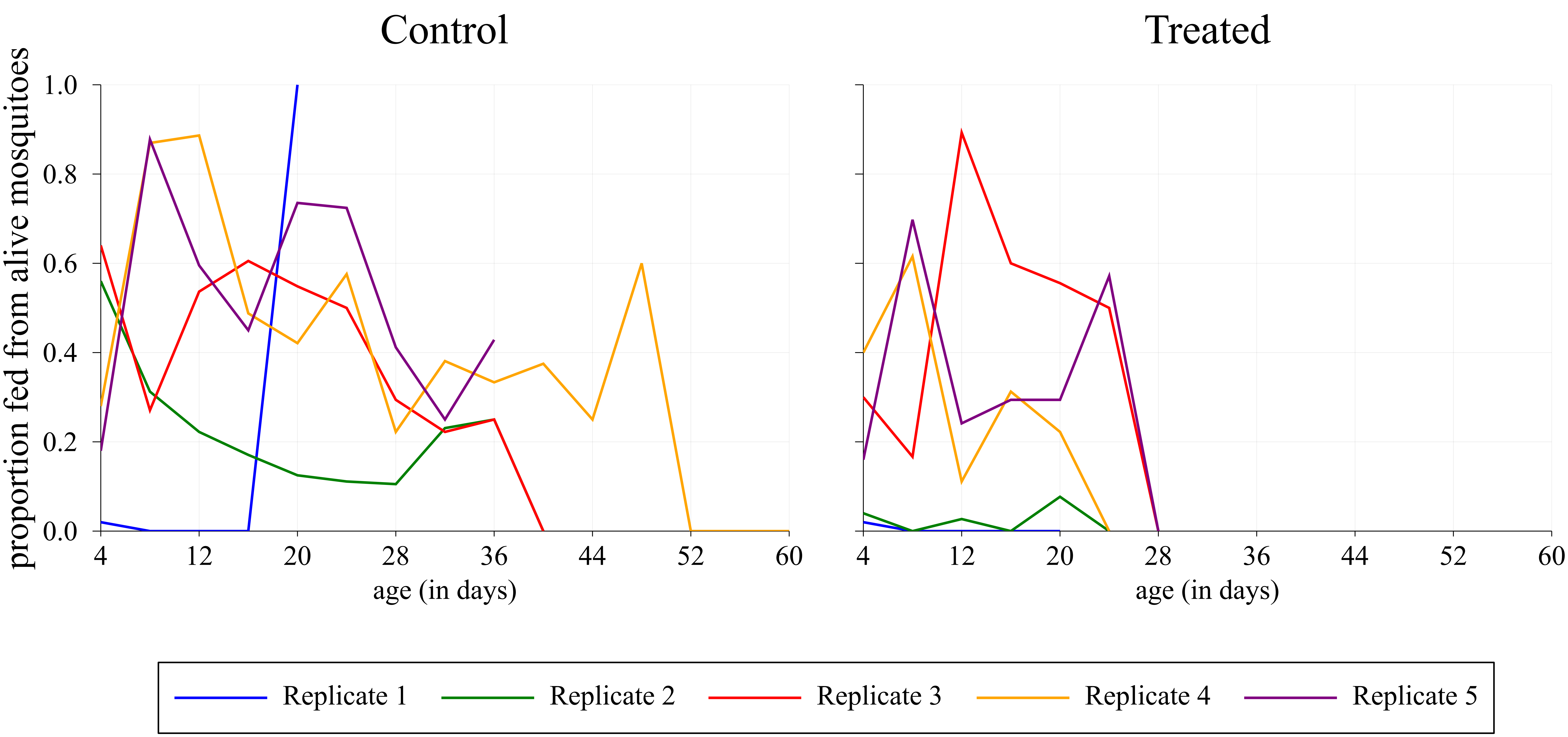

### S2 Fig

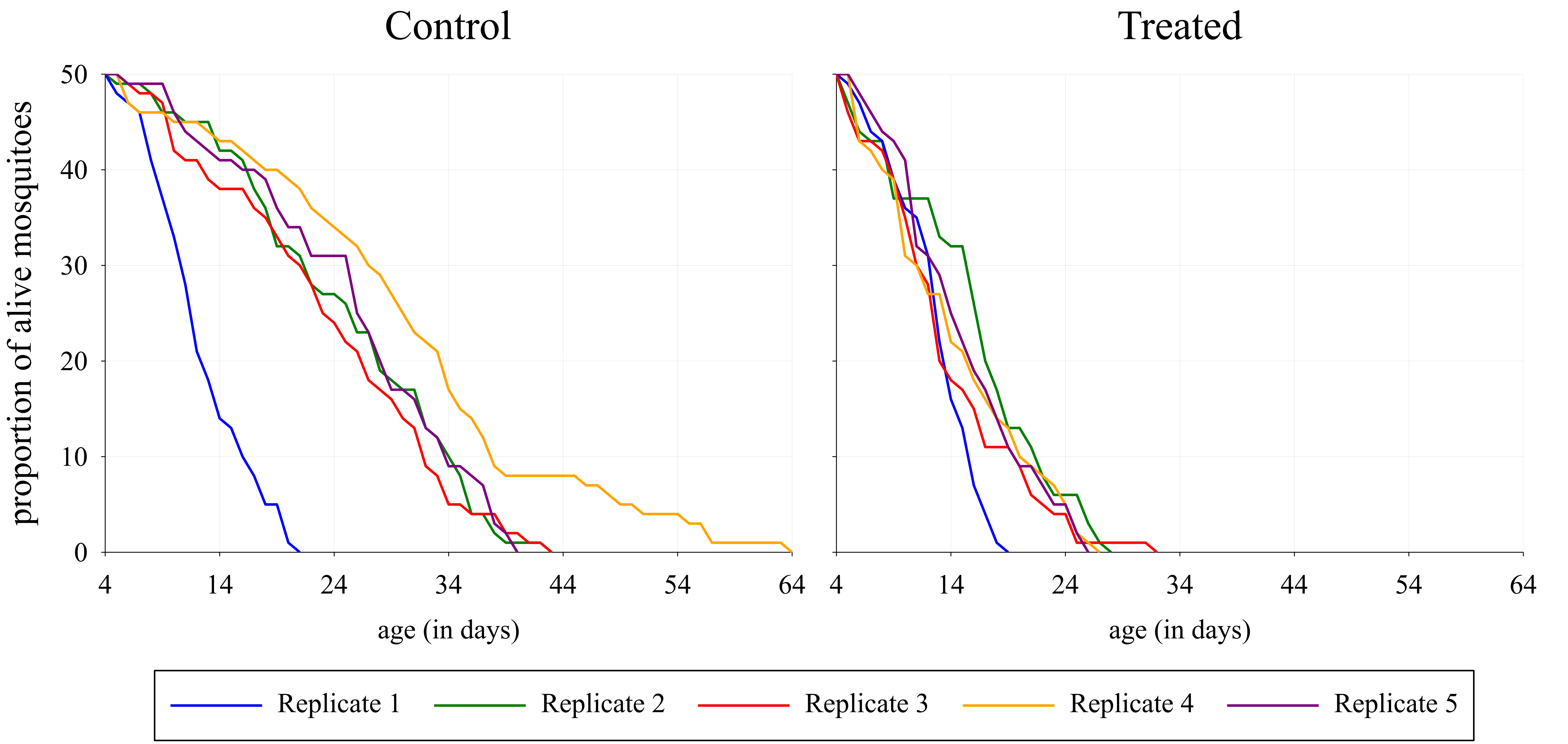

### S3 Fig

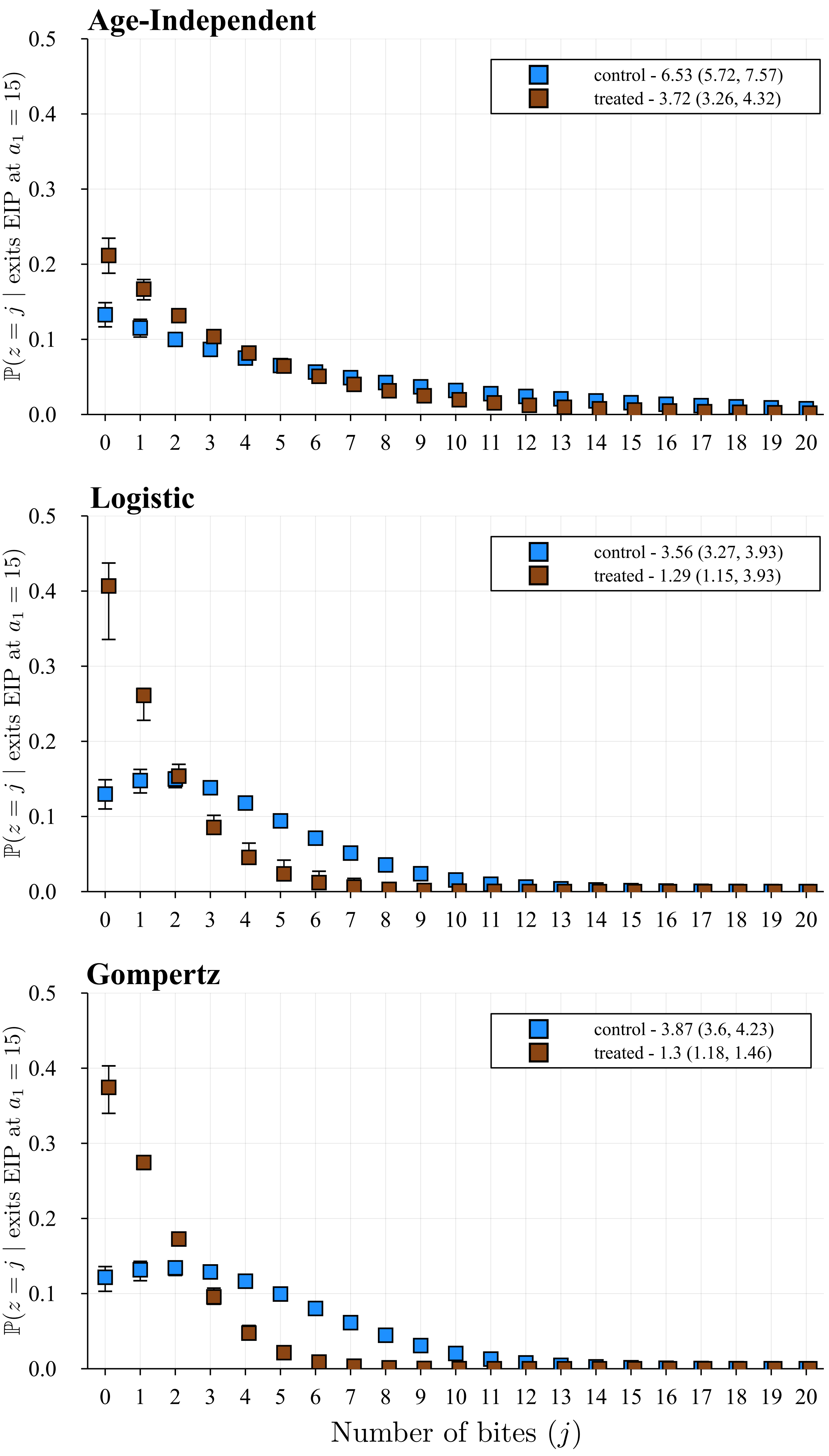
