## Supplementary material for "Omitting age-dependent mosquito mortality in malaria models underestimates the effectiveness of insecticide-treated nets": S1 File

### S1 File: Maximum likelihood estimation and Monte Carlo methods

Melissa A. Iacovidou, Priscille Barreaux, Simon E. F. Spencer, Matthew B. Thomas, Erin E. Gorsich, Kat S. Rock

### Parameter estimation and propagation of uncertainty

The parameter estimations for the various survival functions in the paper were obtained through the maximum likelihood estimation (MLE) method. Using the asymptotic normality of the MLE, we were able to approximate the joint distribution of the parameters. We then took a random sample (of size 10,000) from this distribution in order to perform Monte Carlo simulations for the survival and mortality functions, and the four steps in the calculation of the vectorial capacity. Using the simulations we obtained a 95% confidence region for every result. Here, we outline these methods for clarity.

### Maximum likelihood estimation

In order to use the MLE method, we need the likelihood functions for each of our cases. It is quite common to take the natural logarithm of the likelihood function (log-likelihood), as it is more convenient to work with. By maximising the log-likelihood (or minimising the negative log-likelihood), parameter estimates are obtained. We outline this method for the three considered survival functions.

#### Age-Independent

In the age-independent case we are dealing with the exponential distribution. We have the constant mortality function, $\mu_{AI}$, (which is often referred to as the hazard function in survival analysis) and the survival function obtained from that, $S_{AI}$. We multiply both to obtain the probability density function (PDF) which we need for the MLE:

$$f_{AI}(t;\mu_{\text{const}}) = \mu_{AI}(t)S_{AI}(t) = \mu_{\text{const}}e^{-t\mu_{\text{const}}}.$$

The likelihood function is then defined as:

$$L_{AI}(\mu_{\text{const}};\mathbf{t}) = \prod_{i=1}^{k} f_{AI}(t_i;\mu_{\text{const}}) = \prod_{i=1}^{k} \mu_{\text{const}}e^{-t_i\mu_{\text{const}}}$$

where $k$ is the number of death observations we have in our data set, and $t_i$, with $i = 1,\ldots,k$, are the observed times where each mosquito died. Hence, we obtain the log-likelihood function:

$$\begin{aligned}\ell_{AI}(\mu_{\text{const}};\mathbf{t}) &= \log(L_{AI}(\mu_{\text{const}};\mathbf{t}))\\ &= \sum_{i=1}^{k}\log(f_{AI}(t_i;\mu_{\text{const}}))\\ &= k\log(\mu_{\text{const}}) - \mu_{\text{const}}\sum_{i=1}^{k} t_i\end{aligned}$$

In the exponential case, it is easy to find the value of $\mu_{\text{const}}$ where this is maximum. We simply take the derivative of $\ell_{AI}(\mu_{\text{const}};\mathbf{t})$ with respect to $\mu_{\text{const}}$ and set that to zero $\left(\frac{\mathrm{d}\ell_{AI}}{\mathrm{d}\mu_{\text{const}}} = 0\right)$. This gives us

$$\hat{\mu}_{\text{const}} = \frac{k}{\sum_{i=1}^{k} t_i}.$$

#### Logistic

In the logistic case, where the mortality function ($\mu_L$) is dependent on three parameters, we calculate our PDF using the same method as above, and use it to get the likelihood ($L_L$) and log-likelihood ($\ell_L$) functions:

$$
\begin{aligned}
f_L(t;\mu_1,\mu_2,\mu_3) &= \mu_1 e^{-\mu_1 t}\frac{\left(e^{\mu_2(\mu_3-t)}+1\right)^{-\frac{\mu_1}{\mu_2}-1}}{\left(e^{\mu_2\mu_3}+1\right)^{-\frac{\mu_1}{\mu_2}}} \\
L_L(\mu_1,\mu_2,\mu_3;\mathbf{t}) &= \prod_{i=1}^{k} f_L(t_i;\mu_1,\mu_2,\mu_3) \\
\ell_L(\mu_1,\mu_2,\mu_3;\mathbf{t}) &= \sum_{i=1}^{k}\log(f_L(t_i;\mu_1,\mu_2,\mu_3)) \\
&= k\log(\mu_1) + k\frac{\mu_1}{\mu_2}\log(1+e^{\mu_2\mu_3}) - \mu_1\sum_{i=1}^{k} t_i - \left(\frac{\mu_1}{\mu_2}+1\right)\sum_{i=1}^{k}\log\left(1+e^{\mu_2(\mu_3-t_i)}\right)
\end{aligned}
$$

We can get three equations for our three unknowns by setting the partial derivatives to zero, i.e. $\frac{\partial \ell_L}{\partial \mu_1}=0$, $\frac{\partial \ell_L}{\partial \mu_2}=0$, $\frac{\partial \ell_L}{\partial \mu_3}=0$. However, the values for the parameters have to be computed numerically. Therefore, we use the Optim.jl package in Julia, and we minimise the negative log-likelihood to obtain the estimated values.

Upon sampling using the Monte Carlo method (explained further on), we notice that it is possible to obtain negative samples for $\mu_1$ and $\mu_2$. Hence, we decide to use a transformation for the parameters. We use $\phi(\mu_i)=\log(\mu_i)$ for all three parameters ($i\in\{1,2,3\}$). Therefore, the log-likelihood is as follows:

$$
\ell_L(\phi_1,\phi_2,\phi_3;\mathbf{t}) = k\phi_1 + ke^{\phi_1-\phi_2}\log\left(1+e^{e^{\phi_2+\phi_3}}\right) - e^{\phi_1}\sum_{i=1}^{k} t_i - \left(e^{\phi_1-\phi_2}+1\right)\sum_{i=1}^{k}\log\left(1+e^{e^{\phi_2}(e^{\phi_3}-t_i)}\right)
$$

We use this to sample for $\phi_1$, $\phi_2$, and $\phi_3$, and then transform back to our original parameters, i.e. $\mu_i = e^{\phi_i}$.

#### Gompertz

We follow a similar idea for the Gompertz mortality function ($\mu_G$) as seen above with the logistic case. Hence, we have the following functions for our PDF, likelihood, and log-likelihood respectively:

$$
\begin{aligned}
f_G(t;g_1,g_2) &= g_1\exp\left[\frac{g_1}{g_2}-\frac{g_1}{g_2}e^{g_2 t}+g_2 t\right] \\
L_G(g_1,g_2;\mathbf{t}) &= \prod_{i=1}^{k} f_G(t_i;g_1,g_2) \\
\ell_G(g_1,g_2;\mathbf{t}) &= \sum_{i=1}^{k}\log(f_G(t_i;g_1,g_2)) \\
&= k\log(g_1) + k\frac{g_1}{g_2} + g_2\sum_{i=1}^{k} t_i - \frac{g_1}{g_2}\sum_{i=1}^{k} e^{g_2 t_i}
\end{aligned}
$$

We set the partial derivatives to zero, as before, $\frac{\partial \ell_G}{\partial g_1}=0$, $\frac{\partial \ell_G}{\partial g_2}=0$. However, as with the logistic case, we minimise the negative log-likelihood numerically using Julia to obtain the parameter estimates.

### Wald confidence intervals

We calculate Wald confidence intervals for all the parameters obtained through the MLE method. This is easier to calculate for a single parameter (for example in the age-independent case). Here, the asymptotic normal confidence interval for our parameter $\mu_{\text{const}}$ has the form

$$\hat{\mu}_{\text{const}} \pm z \frac{1}{\sqrt{-\ell''_{AI}(\hat{\mu}_{\text{const}}; \mathbf{t})}},$$

where $z$ is the confidence value, and $\ell''_{AI}(\hat{\mu}_{\text{const}}; \mathbf{t})$ is the second derivative of the log-likelihood function with respect to $\mu_{\text{const}}$, evaluated at $\mu_{\text{const}} = \hat{\mu}_{\text{const}}$. The standard error of $\hat{\mu}_{\text{const}}$ is estimated by $\mathcal{I}(\hat{\mu}_{\text{const}}) = \frac{1}{\sqrt{-\ell''_{AI}(\hat{\mu}_{\text{const}}; \mathbf{t})}}$, the observed Fisher information. The 95% confidence interval of our parameter estimate can be written as

$$\hat{\mu}_{\text{const}} \pm 1.96 \frac{1}{\sqrt{\mathcal{I}(\hat{\mu}_{\text{const}})}}.$$

In our two (Gomperz) or three (logistic) parameter cases we use the Fisher Information matrix (FIM). A typical element of the $N \times N$ FIM is:

$$[\mathcal{I}(\boldsymbol{\theta})]_{ij} = -\mathbb{E}\left[\frac{\partial^2}{\partial\theta_i \partial\theta_j} \ell(\boldsymbol{\theta}; \mathbf{t})\right]$$

where we have the negative expectation of the second derivative of the log-likelihood function with respect to $\theta_i$ and $\theta_j$. $\boldsymbol{\theta}$ is our $N \times 1$ parameter vector $\boldsymbol{\theta} = [\theta_1 \ \theta_2 \ \ldots \ \theta_N]^T$. So the confidence interval for the $i^{\text{th}}$ component of $\boldsymbol{\theta}$ has the form

$$\hat{\theta}_i \pm z\sqrt{(\mathcal{I}(\hat{\boldsymbol{\theta}})^{-1})_{ii}}.$$

### Monte Carlo simulations

We also require confidence intervals of functions of the MLE, and these functions can only be evaluated via numerical integration, we cannot use the standard approach for the propagation of uncertainty. Instead, we draw Monte Carlo samples from the asymptotic distribution of the MLE. We then calculate the appropriate functions for each Monte Carlo sample and use the appropriate quantiles of the resulting Monte Carlo distribution to provide the required confidence intervals. Specifically,

$$\hat{\boldsymbol{\theta}} \sim \mathcal{N}(\boldsymbol{\theta_0}, \mathcal{I}(\boldsymbol{\theta_0})^{-1})$$

where $\boldsymbol{\theta_0}$ is the true value of the parameter vector, and the inverse of the FIM is an estimator of the asymptotic covariance matrix.

#### Bibliography


Agresti, A., 2013. *Categorical data analysis.* 3rd ed. John Wiley & Sons.

Harrison, R.L., 2010, January. Introduction to monte carlo simulation. In *AIP conference proceedings* (Vol. 1204, No. 1, pp. 17-21). American Institute of Physics.

Lehmann, E.L., 1999, *Elements of Large-Sample Theory.* Springer.

Machin, D., Cheung, Y.B. and Parmar, M., 2006. *Survival analysis: a practical approach.* 2nd ed. John Wiley & Sons.

Raychaudhuri, S., 2008, December. Introduction to monte carlo simulation. In 2008 *Winter simulation conference* (pp. 91-100). IEEE.
