## Supplementary material for "Omitting age-dependent mosquito mortality in malaria models underestimates the effectiveness of insecticide-treated nets": S2 File

### S2 File: Rethinking the vectorial capacity – detailed calculations

Melissa A. Iacovidou, Priscille Barreaux, Simon E. F. Spencer, Matthew B. Thomas, Erin E. Gorsich, Kat S. Rock

### Detailed steps to compute expected number of infectious bites

We give more detailed calculations for the four steps regarding the computation of the expected number of infectious bites a mosquito takes in its lifetime. In what follows we investigate three cases; CASE (I) being the age-independent, CASE (II) the logistic, and CASE (III) the Gompertz. As a reminder, these are the mortality functions for the three cases that we are interested in:

$$\begin{aligned} \text{Age-Independent:} \quad & \mu_{AI}(a) = \mu_{\text{const}} \\ \text{Logistic:} \quad & \mu_L(a) = \frac{\mu_1}{1 + e^{\mu_2(-a+\mu_3)}} \\ \text{Gompertz:} \quad & \mu_G(a) = g_1 e^{a g_2} \end{aligned}$$

#### Step 1: $\mathbb{P}$(mosquito survives EIP | infectious blood-meal at $a_0$)

Assuming that the EIP follows an Erlang distribution, i.e.

$$f(t; k, \sigma) = \frac{(k\sigma)^k t^{k-1} e^{-k\sigma t}}{(k-1)!}, \tag{1}$$

we can calculate this probability for the three different cases:

$$\begin{aligned} &\mathbb{P}(\text{surviving EIP} \mid \text{infectious blood-meal taken at age } a_0) \\ &\quad = \int_{t=0}^{\infty} \mathbb{P}(\text{exits EIP after time } t \mid \text{survives to time } t) \times \mathbb{P}(\text{survives from } a_0 \text{ to time } t)\, \mathrm{d}t \\ &\quad = \int_{t=0}^{\infty} [\text{PDF of EIP}(t)] \times \exp\left(-\int_{a_0}^{a_0+t} \mu(x)\,\mathrm{d}x\right)\, \mathrm{d}t \\ &\text{Using Eq (1):} \\ &\quad = \int_{t=0}^{\infty} \frac{(k\sigma)^k t^{k-1} e^{-k\sigma t}}{(k-1)!} \times \exp\left(-\int_{a_0}^{a_0+t} \mu(x)\,\mathrm{d}x\right)\, \mathrm{d}t \end{aligned}$$

CASE (I): $\mathbb{P}$(surviving EIP | infectious blood-meal at age $a_0$)

$$= \int_{t=0}^{\infty} \frac{(k\sigma)^k t^{k-1} e^{-k\sigma t}}{(k-1)!} \times e^{-\mu_{\text{const}} t}\, \mathrm{d}t = \left(\frac{k\sigma}{k\sigma + \mu_{\text{const}}}\right)^k \tag{2}$$

CASE (II): $\mathbb{P}$(surviving EIP | infectious blood-meal at age $a_0$)

$$= \int_{t=0}^{\infty} \frac{(k\sigma)^k t^{k-1} e^{-k\sigma t}}{(k-1)!} \times e^{-\mu_1 t} \left\{\frac{e^{\mu_2(\mu_3-(a_0+t))} + 1}{e^{\mu_2(\mu_3-a_0)} + 1}\right\}^{-\frac{\mu_1}{\mu_2}} \mathrm{d}t \tag{3}$$

CASE (III): $\mathbb{P}$(surviving EIP | infectious blood-meal at age $a_0$)

$$= \int_{t=0}^{\infty} \frac{(k\sigma)^k t^{k-1} e^{-k\sigma t}}{(k-1)!} \times e^{-\frac{g_1}{g_2} e^{a_0 g_2}(e^{g_2 t}-1)}\, \mathrm{d}t \tag{4}$$

Cases (ii) and (iii) cannot be solved analytically, but are integrated numerically using the QuadGK.jl package in Julia.

#### Step 2: $\mathbb{P}(z = j \mid \text{mosquito exits EIP at } a_1)$

We are interested in the probability mass function (PMF) of the number of bites, $z$, supposing the mosquito exits the EIP at age $a_1$, and dies at age $a_2$. This is given by:

$$\mathbb{P}(z = j \mid \text{exits EIP at age } a_1) = \int_{a_1}^{\infty} \mathbb{P}(z = j \mid \text{survive from } a_1 \text{ to } a_2) \times \mathbb{P}(\text{dying at } a_2)\ \mathrm{d}a_2$$

If we assume that the time between bites are exponentially distributed, the first part is given by a Poisson process of the form:

$$\frac{[\alpha(a_2 - a_1)]^j}{j!} e^{-\alpha(a_2 - a_1)} \tag{5}$$

and so it remains to calculate the second part. First, we know that the probability of the mosquito surviving between $a_1$ and $a'$ is equal to $\exp\left(-\int_{a_1}^{a'} \mu(x)\ \mathrm{d}x\right)$ (given our assumption it already survived to age $a_1$). This is equivalent to the probability that the time of death of the mosquito is greater than $a'$, $\mathbb{P}(a_2 > a')$. Hence we can write down the cumulative distribution function of the probability the mosquito dies before age $a'$:

$$F_{a_2}(a') = \mathbb{P}(a_2 \leq a') = 1 - \exp\left(-\int_{a_1}^{a'} \mu(x)\ \mathrm{d}x\right)$$

and so its derivative with respect to $a'$ gives us what we require:

$$f_{a_2}(a') = \frac{\mathrm{d}F_{a_2}(a')}{\mathrm{d}a'} = \mu(a') \exp\left(-\int_{a_1}^{a'} \mu(x)\ \mathrm{d}x\right). \tag{6}$$

Which is the probability density function. We can think of $f_{a_2}(a')\ \mathrm{d}a'$ as the probability of $a_2$ being in the infinitesimal interval $(a', a' + \mathrm{d}a')$. Therefore, using Eqs (5) and (6),

$$\mathbb{P}(z = j \mid \text{exits EIP at age } a_1) = \int_{a_2 = a_1}^{\infty} \frac{\alpha^j (a_2 - a_1)^j}{j!} e^{-\alpha(a_2 - a_1)} \mu(a_2) \exp\left(-\int_{a_1}^{a_2} \mu(x)\ \mathrm{d}x\right)\ \mathrm{d}a_2$$

$$\text{Case (i):} \quad \mathbb{P}(z = j \mid \text{exits EIP at age } a_1) = \frac{\mu_{\text{const}}\alpha^j}{(\mu_{\text{const}} + \alpha)^{j+1}} \tag{7}$$

$$\text{Case (ii):} \quad \mathbb{P}(z = j \mid \text{exits EIP at age } a_1) = \int_{a_2 = a_1}^{\infty} \frac{\alpha^j (a_2 - a_1)^j}{j!} e^{-(\alpha + \mu_1)(a_2 - a_1)} \frac{\mu_1}{1 + e^{\mu_2(\mu_3 - a_2)}} \left\{\frac{e^{\mu_2(\mu_3 - a_2)} + 1}{e^{\mu_2(\mu_3 - a_1)} + 1}\right\}^{-\frac{\mu_1}{\mu_2}} \mathrm{d}a_2 \tag{8}$$

$$\text{Case (iii):} \quad \mathbb{P}(z = j \mid \text{exits EIP at age } a_1) = \int_{a_2 = a_1}^{\infty} \frac{\alpha^j (a_2 - a_1)^j}{j!} e^{a_2 g_2 - (a_2 - a_1)\alpha} g_1 e^{-\frac{g_1}{g_2}(e^{a_2 g_2} - e^{a_1 g_2})}\ \mathrm{d}a_2 \tag{9}$$

Cases (ii) and (iii) are again solved numerically. We further calculate the average number of bites:

$$\mathbb{E}(z \mid \text{exits EIP at age } a_1) = \sum_{j=0}^{\infty} [j \times \mathbb{P}(z = j \mid \text{exits EIP at age } a_1)].$$

**Step 3: $\mathbb{E}(z \mid$ infectious blood-meal at $a_0)$**

We can now calculate the expected number of infectious bites a mosquito takes, given that it takes an infectious blood-meal at age $a_0$:

$$\mathbb{E}(z \mid \text{blood-meal at } a_0) = \sum_{j=0}^{\infty}[j \times \mathbb{P}(z = j \mid \text{blood-meal at } a_0)] \tag{10}$$

Hence, we need to calculate the PMF of the number of bites, given that an infectious blood-meal is taken at age $a_0$. To do so, we use the results from Step 1 and Step 2.

Case (i): To calculate the probability required, we multiply the results from Eqs (2) and (7), however care must be taken for when $j = 0$, where we need to consider that we definitely have zero bites if the mosquito does not survive the EIP.

$$\mathbb{P}(z = j \mid \text{blood-meal at } a_0) = \begin{cases} \left(\frac{k\sigma}{k\sigma+\mu_{\text{const}}}\right)^k \frac{\mu_{\text{const}}\alpha^j}{(\mu_{\text{const}}+\alpha)^{j+1}} & \text{if } j \neq 0 \\ \left(\frac{k\sigma}{k\sigma+\mu_{\text{const}}}\right)^k \frac{\mu_{\text{const}}}{\mu_{\text{const}}+\alpha} + \left(1 - \left(\frac{k\sigma}{k\sigma+\mu_{\text{const}}}\right)^k\right) & \text{if } j = 0 \end{cases}$$

This gives us

$$\mathbb{E}(z \mid \text{infectious blood-meal at } a_0) = \frac{\alpha}{\mu_{\text{const}}}\left(\frac{k\sigma}{k\sigma + \mu_{\text{const}}}\right)^k. \tag{11}$$

Cases (ii) and (iii):

$$\mathbb{P}(z = j \mid \text{blood-meal at } a_0) = \int_{a_1=a_0}^{\infty} [\text{Eq (8) or Eq (9)}] \times \frac{(k\sigma)^k (a_1 - a_0)^{k-1} e^{-k\sigma(a_1-a_0)}}{(k-1)!} \exp\left(-\int_{a_0}^{a_1} \mu(x)\, \mathrm{d}x\right) \mathrm{d}a_1 \tag{12}$$

where we use Eq (8) for case (ii) and Eq (9) for case (iii). The above is true for $j \neq 0$. When $j = 0$, we must add (1 − Eq (3)) to Eq (12) for case (ii), or (1 − Eq (4)) for case (iii), following the same logic as in case (i). This is again integrated numerically (this time using the Cuba.jl package in Julia) and put into Eq (10) to obtain the required solution.

**Step 4: $\mathbb{E}(z)$**

We can now use the previous steps to calculate the expected number of infectious bites a mosquito will take in its lifetime:

$$\mathbb{E}(z) = \int_{a_0=0}^{\infty} \mathbb{E}(z \mid \text{infectious blood-meal at } a_0) \times \mathbb{P}(\text{infectious blood-meal at } a_0)\, \mathrm{d}a_0$$

Breaking this down, the first part is Eq (10), and, for the second part, we must take into consideration that the probability a mosquito gets infected at age $a_0$ depends on the PDF of blood-meals. Specifically, we have:

$$\mathbb{E}(z) = \int_{a_0=0}^{\infty} \mathbb{E}(z \mid \text{infectious blood-meal at } a_0) \times \alpha e^{-\alpha a_0}\, \mathrm{d}a_0 \tag{13}$$

Case (i):

$$\mathbb{E}(z) = \frac{\alpha}{\mu_{\text{const}}}\left(\frac{k\sigma}{k\sigma + \mu_{\text{const}}}\right)^k$$

which, as expected, is the same as Eq (11) as it does not depend on age. Looking back at the vectorial capacity, $C = \frac{m\alpha^2 e^{-\mu n}}{\mu}$ and $C = \frac{m\alpha^2}{\mu}\frac{\sigma}{\sigma+\mu}$, we can see that this is represented here by $\frac{\alpha}{\mu}e^{-\mu n}$ and $\frac{\alpha}{\mu}\frac{\sigma}{\sigma+\mu}$ respectively. This is because the assumed EIP distribution in each case is different (fixed and exponentially distributed), whereas we have assumed an Erlang distribution.

We solve Eq (13) for cases (ii) and (iii) numerically and obtain a single number for each.
